# Biomechanical Characterization of SARS-CoV-2 Spike RBD and Human ACE2 Protein-Protein Interaction

**DOI:** 10.1101/2020.07.31.230730

**Authors:** W. Cao, C. Dong, S. Kim, D. Hou, W. Tai, L. Du, W. Im, X.F. Zhang

## Abstract

The current COVID-19 pandemic has led to a devastating impact across the world. SARS-CoV-2 (the virus causing COVID-19) is known to use receptor-binding domain (RBD) at viral surface spike (S) protein to interact with the angiotensin-converting enzyme 2 (ACE2) receptor expressed on many human cell types. The RBD–ACE2 interaction is a crucial step to mediate the host cell entry of SARS-CoV-2. Recent studies indicate that the ACE2 interaction with the SARS-CoV-2 S protein has higher affinity than its binding with the structurally identical S protein of SARS-CoV-1, the virus causing the 2002-2004 SARS outbreak. However, the biophysical mechanism behind such binding affinity difference is unclear. This study utilizes a combined single-molecule force spectroscopy and steered molecular dynamics (SMD) simulation approach to quantify the specific interactions between CoV-2 or CoV-1 RBD and ACE2. Depending on the loading rates, the unbinding forces between CoV-2 RBD and ACE2 range from 70 to 110 pN, and are 30-50% higher than those of CoV-1 RBD and ACE2 under similar loading rates. SMD results indicate that CoV-2 RBD interacts with the N-linked glycan on Asn90 of ACE2. This interaction is mostly absent in the CoV-1 RBD–ACE2 complex. During the SMD simulations, the extra RBD-N-glycan interaction contributes to a greater force and prolonged interaction lifetime. The observation is confirmed by our experimental force spectroscopy study. After the removal of N-linked glycans on ACE2, its mechanical binding strength with CoV-2 RBD decreases to a similar level of the CoV-1 RBD–ACE2 interaction. Together, the study uncovers the mechanism behind the difference in ACE2 binding between SARS-CoV-2 and SARS-CoV-1, and could aid in the development of new strategies to block SARS-CoV-2 entry.

**STATEMENT OF SIGNIFICANCE:** This study utilizes a combined single-molecule force spectroscopy and steered molecular dynamics simulation approach to quantify the specific interactions between SARS-CoV-2 or SARS-CoV-1 receptor-binding domain and human ACE2. The study reveals the mechanism behind the difference in ACE2 binding between SARS-CoV-2 and SARS-CoV-1, and could aid in the development of new strategies to block SARS-CoV-2 entry.

## INTRODUCTION

Coronavirus Disease-2019 (COVID-19) is a highly contagious infectious disease caused by Severe Acute Respiratory Syndrome-Coronavirus-2 (SARS-CoV-2) (1). First reported in Wuhan, China in December 2019, COVID-19 has rapidly spread to the entire world and become a devastating pandemic. As of now, there is no FDA-approved antiviral or vaccine for COVID-19.

Coronaviruses (CoVs) are enveloped, positive-sense RNA viruses that belong to the family *Coronaviridae* (2). They are classified in four genera (α, β, γ, and δ). Both SARS-CoV-2 and SARS-CoV-1 (which caused the 2002-2004 outbreak) belong to the β-CoV genus. The genomes of SARS-CoV-2 and SARS-CoV-1 share 76% sequence identity (2). Both genomes encode four structural proteins: spike (S), envelope (E), membrane (M) and nucleocapsid (N). The M protein maintains the viral lipid membrane integrity. The E protein facilitates assembly and release of the virus and the N protein encapsulates and protects the viral genome (3).

The S protein (~ 150 kDa) is a heavily N-linked glycosylated homo-trimer projecting 20 nm from the surface of the CoV (3). The trimeric S glycoprotein is a class I fusion protein and mediates attachment to the host receptor. The S1 portion contains the large receptor-binding domain (RBD), and S2 portion forms the stalk of the spike molecule. The atomic structures of the SARS-CoV-2 S protein in a trimeric form (4) as well as the RBD-receptor complex have been determined (5). These structures are similar to the previously reported structures of SARS-CoV-1 S protein (6–8), indicating that the two proteins might function in a similar fashion.

Angiotensin-converting enzyme 2 (ACE2) is a known receptor for SARS-CoV-1 and SARS-CoV-2 S proteins. The major physiological function of ACE2 is to hydrolyze angiotensin II (a vasoconstrictor) into angiotensin-(1–7) (a vasodilator), and thereby to lower blood pressure (9, 10). ACE2 is a type I transmembrane protein expressed in a wide variety of organs including the lungs, heart, kidneys, and intestine (11, 12). Recent structural studies show that ACE2 is a homodimer with each monomer consisting of an N-terminal peptidase domain, a C-terminal Collectrin-like domain, a single-pass transmembrane region, and a short cytoplasmic region (8). The RBD binding region on ACE2 is located in its N-terminal peptidase domain with major contact regions located in the α1 and α2 helixes, as well as the linker between β3 and β4 strands (8).

The binding interactions between ACE2 and CoV S proteins have been widely studied recently. Although there are variations among different binding assays reported, majority of reports show that a higher binding affinity between ACE2 and SARS-CoV-2 S compared to the binding between ACE2 and SARS-CoV-1 S (13, 14). However, the mechanism behind such difference is still unclear. In addition, little is known about the biomechanical strength of ACE2-S interaction that drives viral adhesion and helps withstand the force exerted during viral entry.

In this work, using atomic force microscopy (AFM)-based single-molecule force spectroscopy, a method where a single bond rupture (i.e., interaction) between two molecules can be measured directly, we have quantified the mechanical strengths between ACE2 and SARS-CoV-1 RBD (shortly RBD^CoV1^) or SARS-CoV-2 RBD (shortly RBD^CoV2^). As AFM can measure forces in the pico-Newton (pN) range, it is possible to detect inter-molecular forces, and allow for weak interactions between tip-bound ligands and surface-bound receptor molecules to be quantified in terms of their affinities and rate constants (15). Furthermore, AFM has been recently adopted by us and others to study the interactions between viruses and host cells (16–18). We also used all-atom steered molecular dynamics (SMD) simulations to pull the RBD^CoV1^-ACE2 or RBD^CoV2^-ACE2 complexes with or without N-glycans. Both AFM and SMD confirmed a stronger force/energy associated with the dissociation of RBD^CoV2^-ACE2 complex, and that this enhanced mechanical strength stems from an additional interaction of RBD^CoV2^ with an N-linked glycan of ACE2 Asn90.

## MATERIALS AND METHODS

### Protein expression

Immortalized HEK 293T cells purchased from American Type Culture Collection (ATCC) were cultured in DMEM medium (ATCC), and supplemented with 4 mM L-glutamine, 4500 mg/L glucose, 1 mM sodium pyruvate, 1500 mg/L sodium bicarbonate, 1% penicillin streptomycin, and 10% fetal bovine serum. RBD proteins were expressed as previously described with some modifications (19). Briefly, genes encoding SARS-CoV-1 RBD and SARS-CoV-2 RBD proteins containing a C-terminal Fc tag were amplified by PCR using codon-optimized SARS-CoV-1 or SARS-CoV-2 S plasmid, and inserted into a human Fc expression vector (Invitrogen). The proteins were expressed in HEK293T cells, and purified by protein A affinity chromatography (GE Healthcare).

### Cantilever preparation/ coverslip preparation

To functionalize AFM cantilevers (MLTC, Bruker Nano) with RBD, the cantilever was first silanized with 3-(trimethoxysilyl)propyl methacrylate to obtain surface thiol groups. RBD^CoV1^, RBD^CoV2^, or Middle East Respiratory Syndrome Coronavirus (RBD^MERS-CoV^, as a negative control) were immobilized onto a (3-aminopropyl)-triethoxysilane sinalized AFM cantilever (MLTC, Bruker Nano) using a heterobifunctional polyethylene glycol (PEG) crosslinker, Acetal-PEG-NHS (Creative PEGworks), according to the protocol developed by Dr. Hermann J. Gruber, Johannes Kepler University (17). Soluble recombinant ACE2 (ACRO Biosystems) was attached to the silanized glass coverslips using the same crosslinking approach. Functionalized cantilevers and glass surfaces were stored in PBS (3 × 5 min) and used for AFM experiment within 8 hours.

### Single-molecule force measurements

All single-molecule force measurements were conducted using a custom-designed AFM apparatus. AFM measurements were collected at cantilever retraction speeds ranging from 0.19 to 7.5 μm/s to achieve the desired loading rate (5,000-20,000 pN/s). All measurements were conducted at 25°C in Phosphate-buffered saline (PBS). The contact time and indentation force between the cantilever and the sample were minimized to obtain measurements of the unitary unbinding force.

To enable measurement of a single molecular interaction, the contact between the cantilever tip and the substrate was minimized by reducing both the contact duration (<50 ms) and the compression force (100-200 pN). The brief contact duration was chosen to ensure that, for the majority of contacts (67% or greater), no adhesion (rupture force) was observed between AFM tip and surface. Assuming the adhesion bond formation obeyed Poisson statistics, an adhesion frequency of ~33% in the force measurements implies that among the observed unbinding events, the probabilities of forming a single, double, and triple adhesion bonds between AFM tip and surface were 81%, 16%, and 2%, respectively (20). Therefore, our experimental condition ensured there was a >80% probability that the adhesion event was mediated by a single bond (21).

### Statistical analysis

For each pulling speed, over 500 force curves were recorded, which yielded 40 to 200 unbinding forces. Curve fitting was performed using IGOR Pro or Origin software by minimizing the chi-square statistic for the optimal fit. Unless otherwise stated, the data is reported as the mean and the standard error of the estimate. Statistical analyses between groups were performed using an unpaired t-test or ANOVA, with a p-value less than 0.05 considered to be statistically significant.

### Steered molecular dynamics simulation

All SMD simulations were performed using NAMD (22) The CHARMM36(m) (23, 24) force field was used for protein and carbohydrates. PDB ID 2AJF (6) from Protein Data Bank was used for an RBD^CoV1^-ACE2 complex structure and PDB ID 6VW1 (5) for an RBD^CoV2^-ACE2 complex structure. We used a TIP3P water model (25), and K^+^ and Cl^-^ ions with a concentration of 0.15 M were added to neutralize the system. All simulation systems and parameters were set up through CHARMM-GUI *Solution Builder* (26, 27). Analysis were done with CHARMM (28) and visualization through VMD (29).

PDB:6VW1 has five N-linked glycans in ACE2 (Asn53, Asn90, Asn103, Asn322, and Asn549) and one N-glycan in RBD^CoV2^ (Asn343), and PDB:2AJF has four N-linked glycans in ACE2 (Asn53, Asn90, Asn322, and Asn549) and one N-glycan in RBD^CoV1^ (Asn330). Similar to other crystal or cryo-EM structures, all N-glycan structures in both PDB structures are incomplete, as they are truncated in experiment or not observable due to low resolution and high structural flexibility. Since we did not know glycoforms of the ACE2 glycosylation sites at the time of this study, we used the N-glycan core pentasaccharide (a minimum structure of all N-glycans: **Fig. S1**) in all N-glycosylation sites including Asn103 of ACE2 in PDB:2AJF. *Glycan Reader & Modeler* (30–32) in CHARMM-GUI was used to model N-glycan core pentasaccharide in all glycosylation sites using the templates from GFDB (33) (Glycan Fragment Database). To compare the receptor-binding affinity between RBD^CoV1^ and RBD^CoV2^ and to explore influences of N-glycans on binding affinity, we made four systems: S^CoV1+G^ (RBD^CoV1^-ACE2 with N-glycans), S^CoV1-G^ (RBD^CoV1^-ACE2 without N-glycans), S^CoV2+G^ (RBD^CoV2^-ACE2 with N-glycans), and S^CoV2-G^ (RBD^CoV2^-ACE2 without N-glycans).

For the SMD simulations, the protein complex structures were initially aligned along the X-axis in a cubic water box with an initial size of 171 Å for S^CoV1±G^ and 172 Å for S^CoV2±G^; a total number of atoms is about 470,000. The pulling forces were applied to the center of mass (COM) of each protein (i.e., RBD and ACE2). In the pulling process, the spring constant was set to 5 kcal/mol/Å^2^ and its moving speed to 0.5 Å/ns in the opposite directions along the X-axis. Gentle restrains with a force constant of 5 kcal/mol/Å^2^ were applied to each protein’s COM to restrict their movement along the Y/Z directions during the pulling process. The SMD simulations stopped at 30 ns when two proteins were detached from each other. 9 independent simulations for each system were performed for better statistics.

The van der Waals interactions were smoothly switched off over 10-12 Å by a force-based switching function (34). The electrostatic interactions were calculated by the particle-mesh Ewald method with a mesh size of 1 Å for fast Fourier transformation and sixth order B-spline interpolation. SHAKE algorithm was used to constrain bond lengths involving hydrogen atom (35) and the simulation time-step was set to 2 fs. We first relaxed the system in an NVT (constant particle number, volume, and temperature) ensemble at 303.15 K with harmonic restraints to all solute atoms. The constant temperature was controlled by Langevin dynamics with a damping frequency of 50 fs^-1^. 100-120 ps NPT (constant particle number, pressure, and temperature) simulation was then applied to adjust the solvent density. The Langevin piston method was used to control the pressure. A dihedral restraint with a force constant of 1 kcal//mol/rad^2^ was applied to carbohydrates to keep the carbohydrate chair conformation during these equilibration steps. To perform the SMD simulation, a COLVARS method was used (36), and the COMs of two proteins were calculated first and used as the external forces’ initial positions. The effective spring potential (whose negative derivative is used to represent external forces) acting on the COM of each protein was calculated using the following equation: 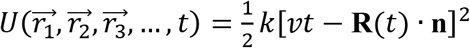, where *k* is the spring constant, *v* is the moving speed of the spring potentials, **R**(*t*) is the current position of the selected protein COM, and **n** is the unit vector along the protein COMs. As a result of this spring potential, the spring-connected protein would move following the energy well, so that two proteins are pulled apart.

## RESULTS

### RBD of SARS-CoV-2 S protein binds ACE2 stronger compared to SARS-CoV-1

First, we characterized the mechanical interaction between the RBD of SARS-CoV S proteins and ACE2 using AFM. We have attached RBD^CoV1^, RBD^CoV2^, or RBD^MERS-CoV^ (negative control) to a micro-cantilever, the force probe, via an established protocol using polyethylene glycol coupling chemistry (17, 18). The cantilever-bound RBD was controlled to interact with surface-immobilized soluble ACE2 via AFM force scans (**Fig. 1A**).

**Fig. 1.**
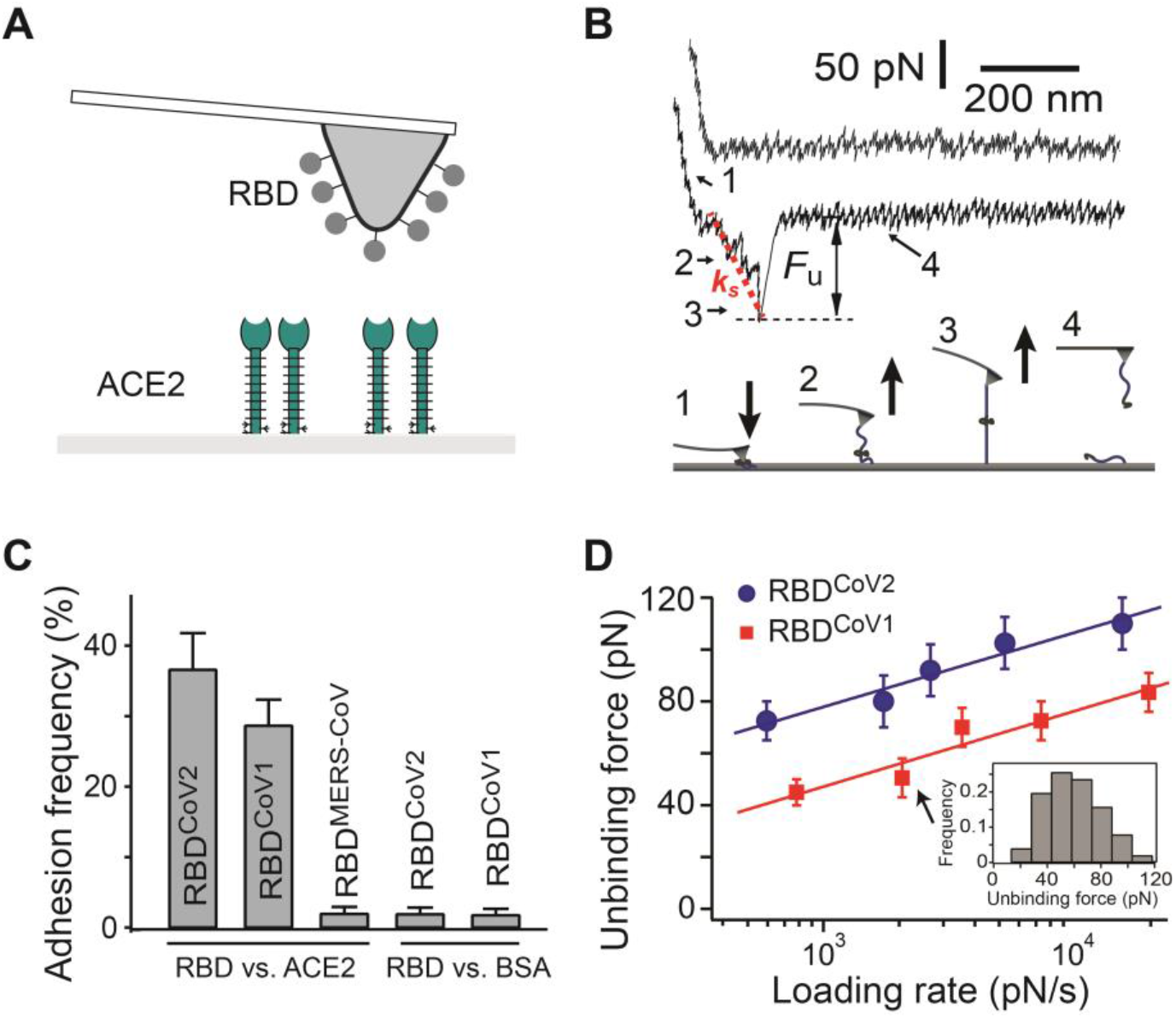
Single-molecule studies of CoV RBD-ACE2 interactions. **(A)** Schematic of the experimental system. The micro-cantilever is functionalized with RBD. Soluble human ACE2 is immobilized on the opposing surface using established protocols. **(B)** The upper panel shows two sample AFM pulling traces of the RBD^CoV2^-ACE2 interaction. The first (upper) trace had no interaction and the second (lower) trace shows the rupture force of the protein-protein complex. *F_u_* is the unbinding force. *k_s_* is the system spring constant that is derived from the slope of the force-displacement trace. *k_s_* is used to derive the loading rate for individual unbinding events. The lower panel illustrates the four stages of stretching and rupturing of a single RBD-ACE2 complex using the AFM. **(C)** Interaction specificity was shown by the adhesion frequency measurement for different interacting pairs. Contact force, contact time, and retraction speed for all the interacting AFM tip and surfaces were set at 150 pN, 0.1 s, and 3.7 μm/s, respectively. Error bars are Poisson errors (i.e., the square root of the adhesion number). **(D)** The dynamic force spectra (i.e., the plot of most probable unbinding force (*F_u_^*^*) as a function of loading rate (*r_f_*) of the RBD-ACE2 interactions. The data is fitted to the single-barrier Bell-Evans model to extract the off-rate k_off_ (37). Inset: a representative histogram to determine the most probable unbinding force.

All single-molecule force measurements were conducted using a custom-built AFM designed for operation in the force spectroscopy mode (38–42). Using a piezoelectric translator, the functionalized cantilever was lowered onto an ACE2-functionalized surface to allow possible binding between RBD and ACE2 to occur. After a brief contact, the cantilever was retracted from the surface. Any binding interaction between tip and substrate would lead to an adhesive pull-off force determined from the deflection of the cantilever via a position-sensitive two-segment photodiode. (**Fig. 1A** and **1B**, lower panel). **Fig. 1B** shows two typical pulling traces. The first (upper) trace represents a majority (65-70%) of all the pulling curves, showing no interaction (i.e., no adhesive force) between the AFM tip and sample surface. The second (lower) trace, representing approximately 30% of the pulling curves in our single-molecule assay, shows the unbinding (i.e., pull off) force of the tip-substrate interaction. The unbinding force (*F_u_*) of the receptor-ligand complex is derived from the force jump that accompanies the unbinding of the complex. *k_s_* is the system spring constant derived from the slope of each pulling trace.

Interaction specificity was shown by the adhesion frequency measurement under the same measurement conditions. **Fig. 1C** shows a significant decrease in adhesion when either RBD^CoV1^, RBD^CoV2^, or ACE2 was absent, confirming that the vast majority of the recorded unbinding forces stemmed from specific interactions. RBD^MERS-CoV^ and BSA were used here as negative control proteins, as RBD^MERS-CoV^ does not bind ACE2 and its known receptor is dipeptidyl peptidase-4 (43).

The biophysical properties of RBD-ACE2 interactions were studied by the means of a dynamic force spectrum (DFS), and the results are shown in **Fig. 1D**. The DFS is the plot of most probable unbinding force as a function of loading rate. The loading rate is obtained by multiplying the system’s spring constant (**Fig. 1B**) and the pulling speed of each force curve. The unbinding forces of each RBD-ACE2 interactions were first grouped into 5 groups by their loading rates. The distribution of forces within the same group was analyzed by histograms (see inset for one example in **Fig. 2D**). The most probable unbinding forces were then determined from the modes of each histograms. **Fig. 2D** shows that the unbinding force of both RBD-ACE2 complexes increased linearly with the logarithm of the loading rate. However, the unbinding forces of RBD^CoV2^–ACE2 are stronger, ranging from 70 to 110 pN over a loading rate of 5,000 to 20,000 pN/s, whereas the RBD^CoV1^–ACE2 unbinding forces are 30-50% lower under similar loading rates.

**Fig. 2.**
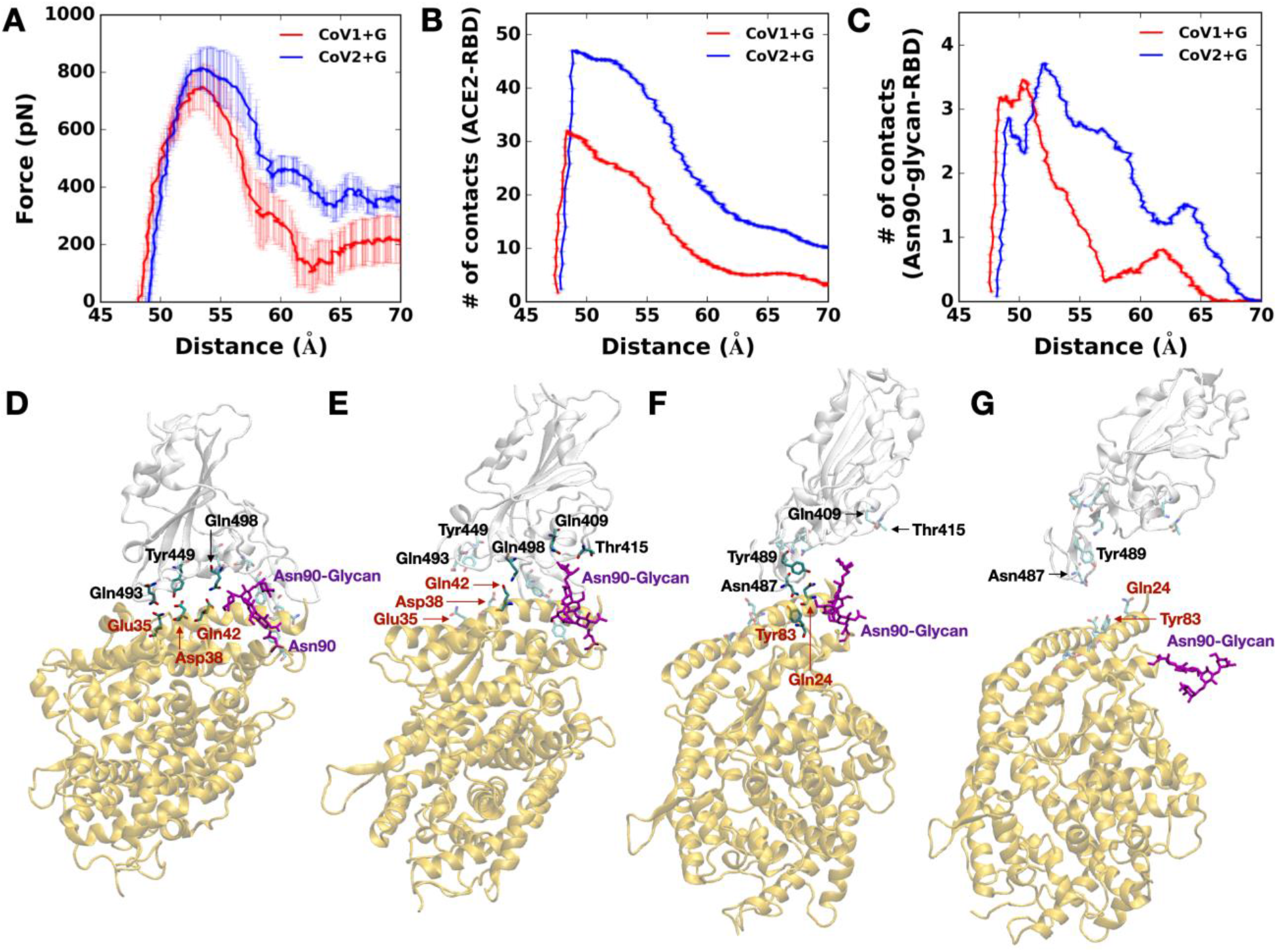
Steered molecular dynamics simulation results. **(A)** Average force profiles of S^CoV1+G^ (red) and S^CoV2+G^ (blue) as a function of distance (*D*^RBD-ACE2^) between the centers of mass of RBD and ACE2. **(B)** Average numbers of contacts between RBD and ACE2 in S^CoV1+G^ (red) and S^CoV2+G^ (blue). **(C)** Average numbers of contacts between RBD and ACE2 Asn90-glycan in S^CoV1+G^ (red) and S^CoV2+G^ (blue). In (A-C), the average data are obtained based on 9 independent SMD simulations for each system, and error bars represent the standard deviations with 68% confident intervals. **(D-G)** Representative snapshots of SMD simulations of S^CoV2+G^ at *D*^RBD-ACE2^ of (D) 49 Å, (E) 57 Å, (F) 65 Å, and (G) 70 Å. Key interacting residues are depicted as the solid sticks and residues losing their interactions are shown as the transparent sticks. The black residue names are for RBD^CoV2^ and brown ones for ACE2. The RBD^CoV2^ and ACE2 are shown by transparent light gray and yellow, respectively. Asn90-glycan is colored in purple.

A more detailed analysis of the biophysical properties of RBD-ACE2 interactions was conducted by fitting the acquired DFS data to the Bell-Evans model. The model describes the influence of an external force on the rate of bond (i.e., complex) dissociation (44). According to this model, a pulling force (*F*) distorts the intermolecular potential of a ligand-receptor complex, leading to a lowering of the activation energy and an increase in the dissociation rate k(f) (or a decrease of bond lifetime t(*F*)) as follows:

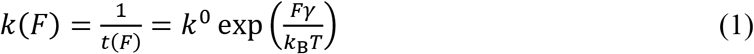

where *k^0^* is the dissociation rate constant in the absence of a pulling force, *γ* is the position of the transition state, T is the absolute temperature, and *k*_B_ is the Boltzmann constant. For a constant loading rate (*r_F_*), the model can be described as:

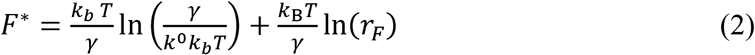

Hence, as predicted by the model, the most probable unbinding force *F*^*^ is a linear function of the logarithm of the loading rate. Experimentally, *F*^*^ was determined from the mode of the unbinding force histograms. Fitting the DFS of RBD^CoV2^–ACE2 interaction to the Bell-Evans model (Eq. 2) yielded a *k^0^* of 0.15 s^-1^, and a *γ* (i.e., activation barrier width) of 0.33 nm. The best-fit parameters for RBD^CoV1^–ACE2 are tabulated in **Table 1**. Clearly, compared to RBD^CoV2^, RBD^CoV1^ binds ACE2 with a 12-fold larger *k^0^* and the same *γ*, indicating that the RBD^CoV2^–ACE2 interaction is stronger with a much slower dissociation rate.

**Table 1.** Bell-Evans model parameters of RBD-ACE2 interactions. Uncertainties are the standard errors of the fits. dgACE2 represents deglycosylated ACE2 treated with PNGase F.

| *Receptor-ligand pairs* | $k^0 \text{ (s}^{-1}\text{)}$ | $\tau^0 \text{ (s)}$ | $\gamma \text{ (nm)}$ |
| --- | --- | --- | --- |
| RBD<sup>CoV2</sup> vs. ACE2 | $0.15 \pm 0.13$ | $6.7 \pm 5.7$ | $0.33 \pm 0.04$ |
| RBD<sup>CoV1</sup> vs. ACE2 | $1.8 \pm 1.3$ | $0.55 \pm 0.40$ | $0.33 \pm 0.06$ |
| RBD<sup>CoV2</sup> vs. dgACE2 | $15.8 \pm 2.4$ | $0.06 \pm 0.01$ | $0.26 \pm 0.02$ |
| RBD<sup>CoV1</sup> vs. dgACE2 | $13.4 \pm 1.6$ | $0.07 \pm 0.01$ | $0.23 \pm 0.01$ |

### SMD identifies an additional interaction of SARS-CoV-2 S RBD with an N-linked glycan of ACE2 Asn90

To gain molecular insight into the receptor-binding affinity between RBD^CoV1^ and RBD^CoV2^ and to explore influences of N-glycans on binding affinity, we performed SMD simulations on the following four systems: S^CoV1+G^ (RBD^CoV1^-ACE2 with N-glycans), S^CoV1-G^ (RBD^CoV1^-ACE2 without N-glycans), S^CoV2+G^ (RBD^CoV2^-ACE2 with N-glycans), and S^CoV2-G^ (RBD^CoV2^-ACE2 without N-glycans). To compare RBD^CoV2^-ACE2 interactions with RBD^CoV1^-ACE2 interactions, pulling force analysis was performed as a function of distance (*D*^RBD-ACE2^) between the COMs of RBD and ACE2. In addition, to investigate how many residues between RBD and ACE2 interact as a function of *D*^RBD-ACE2^, the number of contacts analysis was performed, where a contact was counted if any heavy atom of RBD was within 4.5 Å from any heavy atom of ACE2.

As shown in **Fig. 2A**, the overall force profile of S^CoV2+G^ shows higher forces than s^CoV1+G^ due to greater numbers of RBD^CoV2^-ACE2 contacts compared to RBD^CoV1^-ACE2 (**Fig. 2B**). Initially, s^CoV2+G^ has more contacts than S^CoV1+G^ and the difference in the number of contacts between S^CoV2+G^ and S^CoV1+G^ is about 20 at *D*^RBD-ACE2^ of 52 Å (**Fig. 2B**). The difference decreases to about 17 starting from 55 Å and to 9 at 65 Å, where ACE2 Asn90-glycan maintains its interactions with RBD^CoV2^, whereas such interactions are lost in S^CoV1+G^ (**Fig. 2C**). Note that the force profile in S^CoV2+G^ has a plateau around 60 Å and a small peak around 66 Å, which are attributed to the interactions between ACE2 Asn90-glycan and RBD^CoV2^ from 55 Å to 65 Å (**Fig. 2C**). Because of relatively negligible interactions between ACE2 Asn90-glycan and RBD^CoV1^, the plateau is not observed around 60 Å in S^CoV1+G^. This indicates that the interaction between ACE2 Asn90-glycan and RBD^CoV2^ somewhat blocks the direct contact between RBD^CoV2^ and ACE2 at 55 Å < *D*^RBD-ACE2^ < 65 Å, suggesting that ACE2 Asn90-glycan can hinder the association of RBD^CoV2^ to ACE2 more than RBD^CoV1^, but makes RBD^CoV2^-ACE2 dissociation harder than RBD^CoV1^-ACE2.

Using S^CoV2+G^ as an example, the overall RBD and ACE2 dissociation during the pulling simulation can be divided into three states: state I (<55 Å, **Fig. 2D**), state II (56~70Å, **Fig. 2E, F**), and state III (>70 Å, **Fig. 2G**). In state I, RBD^CoV2^-ACE2 has a number of interactions. As *D*^RBD-ACE2^ increases to 56 Å (state II), RBD^CoV2^ and ACE2 start to lose some of its polar interactions (RBD^CoV2^-ACE2: Gln493-Glu35 and Tyr449-Asp38), but the interaction between Gln498 and Glu42 is intact. Note that ACE2 Asn90-glycan has polar interactions with Gln409 and Thr415 (**Fig. 2E**). At 65 Å (**Fig. 2F**), Asn487 and Try489 of the RBD^CoV2^ loop can still interact with ACE2 Tyr83 and Gln24 due to flexibility of the loop, and Asn487 can also contact Gln24 time to time. At this period, Asn90-glycan loses its contacts with RBD^CoV2^. In state III, RBD^CoV2^ and ACE2 are fully detached with no close interactions (**Fig. 2G**). While the average forces show a subtle difference in between S^CoV1+G^ and S^CoV1-G^ when RBD^CoV1^ and ACE2 start to detach at *D*^RBD-ACE2^ = 56 Å (**Fig. S2A**), S^CoV2+G^ clearly has higher forces over 56 Å to 70 Å than s^CoV2-G^ (**Fig. S2B**). And, RBD^CoV2^ shows slightly higher forces than RBD^CoV1^ even with no glycans, although the differences are within the error bars (**Fig. S2C**).

### Removal of ACE2 N-linked glycans leads to a decrease of unbinding forces

In light of the SMD results, we tested the effect of ACE2 N-linked glycan on the mechanical strength of RBD-ACE2 interactions. To remove the ACE2 N-linked glycans, surface-immobilized ACE2 was incubated with PNGase F (New England Biolabs) for one hour at 37 °C. The effect of PNGase F treatment was analyzed by SDS-PAGE (**Fig. 3A**). After one hour of treatment, the molecular weight of ACE2 was visibly reduced from approximately 115 to 95 kDa. Assuming each N-linked glycosylation adds 2.5 kDa of molecular mass, the result is consistent with seven N-glycosylation sites on ACE2.

**Fig. 3.**
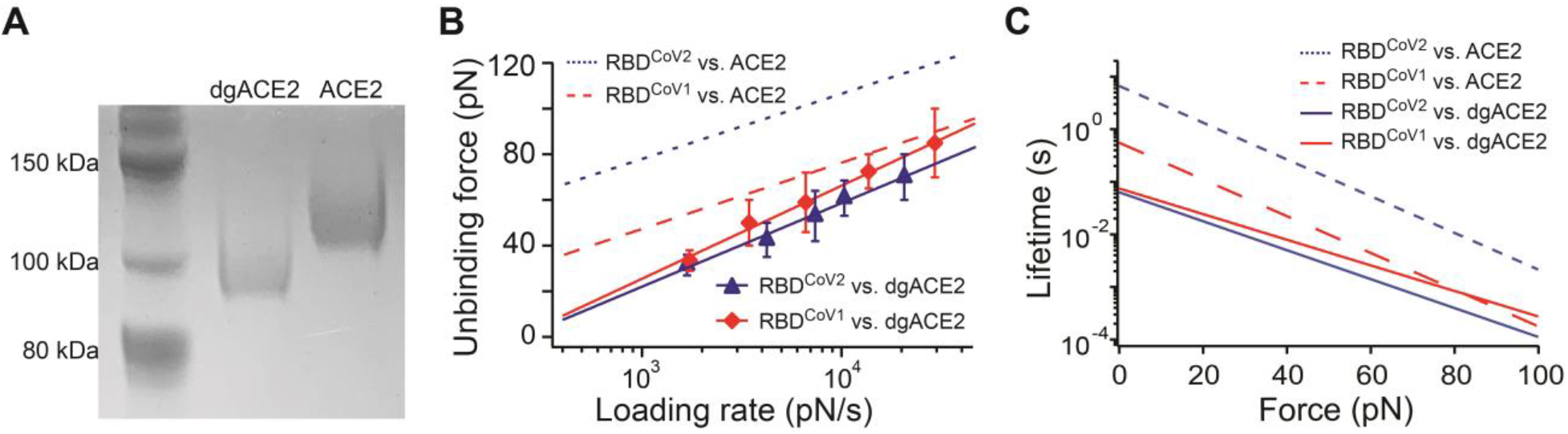
Effect of ACE2 N-glycans on RBD-ACE2 interaction. (A) Deglycosylation (dgACE2) was performed by treatment with PNGase F for one hour at 37°C. Deglycosylation was confirmed via SDS-PAGE stained with EZBlue. (B) The dynamic force spectra of the RBD–dgACE2 interactions. Solid lines are linear fits to Eq. 2 for the interactions. Dashed line is the linear fits for N-glycosylated ACE2 interactions taken from **Fig. 1**. (C) Comparison of lifetimes of RBD-ACE2 complex as a function of force.

Next, AFM unbinding experiments were performed between tip-immobilized RBD and surface immobilized, PNGase F-treated ACE2. As shown in **Fig. 3B**, N-linked glycan removal resulted in a significant decrease of the unbinding forces between RBD^CoV2^ and ACE2, from 70-110 pN to 30-60 pN. The unbinding forces between RBD^CoV1^ and ACE2 also decreased, but to a lesser extent. The DFS of RBD^CoV2^ and RBD^CoV1^ are almost overlapped with each other. This trend is also similar to the SMD results, showing that the force profiles of S^CoV1-G^ and S^CoV1-G^ are within the error bars t (**Fig. S2C**). The Bell-Evans model fit confirmed that after N-glycan removal, the k^0^ of RBD^CoV2^–ACE2 interaction increase by 105 fold (from 0.15 s^-1^ to 15.8 s^-1^), whereas the k^0^ of RBD^CoV1^–ACE2 interaction increase by only 7 fold.

## DISCUSSION AND CONCLUSIONS

Interactions between viral protein and host receptors require direct physical contact between viral and host cell membranes. Unlike interactions in solution (3D), which have at least one interacting molecular species in the fluid phase, the interactions between receptors and ligands anchored on two opposing membranes (2D) are constrained in molecular movement or transport and are under common tensile force. Hence, the 2D reaction kinetics are oftentimes different from 3D kinetics (45, 46). In order to study the mechanism underlying virus-cell interaction, therefore, it is necessary to probe the interaction between anchored molecules using 2D binding assays such as the single-molecule AFM used in this study. Using this method, we found that the dissociate rate for RBD^CoV2^–ACE2 and RBD^CoV1^–ACE2 bonds (or interactions) are significantly different. The one order magnitude slower dissociation rate could partially account for the greater infectivity of SARS-CoV-2.

The Bell-Evans model allows estimation of bond lifetime at different constant pulling forces. Taking the lifetimes and barrier position parameters from **Table 1**, we compared the lifetime time of RBD-ACE2 bonds as a function of force (**Fig. 3C**). At no force, the lifetime of a RBD^CoV2^–ACE2 bond is estimated to be 6.7 s, and is approximately one order of magnitude longer than the lifetime of a RBD^CoV1^–ACE2 bond. Under pulling forces, the lifetimes of both bonds decrease exponentially with force, though the ~10 fold difference between the two bonds remains. This indicates that compared to SARS-CoV-1, SARS-CoV-2 can stay much longer on an ACE2 expressing surface due to stronger RBD-ACE2 binding. After N-glycan removal on ACE2, the unstressed lifetime of both RBD^CoV2^–ACE2 and RBD^CoV2^–ACE2 bonds decreases to 0.06-0.07 s, suggesting that N-glycans may be required for stable SARS-CoV-2 binding to ACE2 much more than SARS-CoV-1.

Using the *k^0^* values from **Table 1**, we were able to estimate the activation energy differences among different scenarios. Estimates of the energy difference between the transition states were calculated as ΔG_12_ = *k*_B_T ln(k_1_/k_2_) where k_1_ and k_2_ are the dissociation rate constants of transition of two interactions used for comparison, respectively. Using this equation, the activation energy barrier for RBD^CoV1^–ACE2 bond dissociation is estimated to be 2.5 *k*_B_T lower than that of the RBD^CoV2^–ACE2 bond. After deglycosylation of ACE2, the activation barrier heights are lower by 4.6 (RBD^CoV2^) and 2 *k*_B_T (RBD^CoV1^), compared to the binding of glycosylated ACE2.

SMD simulations provide molecular-level insight into RBD^CoV^-ACE2 interactions and help us to interpret the AFM data. The SMD simulations manifest that RBD^CoV2^ interacts stronger with ACE2 than RBD^CoV1^ because the former has more direct contacts with ACE2 than the latter. In particular, ACE2 Asn90-glycan appears to have an important role in having stronger interactions with RBD^CoV2^ than RBD^CoV1^ by retaining contacts with residues of RBD^CoV2^, Gln409 and Thr415, even when the original contacts of RBD^CoV2^-ACE2 start to lose (**Fig. 2E**). This additional interaction implies that ACE2 Asn90-glycan can have effects on association and dissociation of RBD^CoV2^-ACE2. In other words, ACE2 Asn90-glycan could hinder the association of RBD^CoV2^ with ACE2 more than RBD^CoV1^, but make RBD^CoV2^-ACE2 dissociation harder than RBD^CoV1^-ACE2. In addition, based on the SMD simulations, we propose a three-step dissociation mechanism of RBD^CoV2^-ACE2 complex.

It should be noted that the current models utilize only RBD out of trimeric SARS-CoV-2 S protein and the N-glycan core structure for all N-glycans. Having a fully-glycosylated SARS-CoV-2 S protein and ACE2 models would provide further insight into the RBD-ACE2 interactions. With a recently modeled fully glycosylated SARS-CoV-2 S protein model (47) and recently-determined glycosylation patterns of ACE2 (48), we plan to study the RBD-ACE2 interactions in a more realistic model.

In conclusion, the study shows the biomechanical parameters important for CoV to attach to host cells. Our results reveal important viral–host cell interaction through ACE2 Asn90-glycan, which could be a potential target for antiviral intervention.

## ACKNOWLEDGEMENTS

This work was supported in part by an NIH grant R15AI133634 and an NSF grant 1804117 (to X.F.Z.), NIH GM126140, NSF DBI-1707207, and MCB-1810695 (to W.I.), NIH grant R01AI157975 (to L.D.), and an internal grant from Lehigh University to X.F.Z, and W.I.

## AUTHOR CONTRIBUTIONS

W.C. designed study, performed experiments, and analyzed results. C.D. and S.K. designed study, performed simulations, analyzed results and wrote the manuscript. D.H. and W.T. performed experiments. L.D. analyzed results and wrote the manuscript. X.F.Z. and W.I. designed study, analyzed results and wrote the manuscript.

